# Neural correlates of orbital telorism

**DOI:** 10.1101/2021.01.07.425611

**Authors:** Mikolaj A. Pawlak, Maria J. Knol, Meike W. Vernooij, M. Arfan Ikram, Hieab H.H. Adams, T. E. Evans

## Abstract

Orbital telorism, the interocular distance, is a clinically informative and in extremes is considered a minor physical anomaly. While its extremes, hypo- and hypertelorism, have been linked to disorders often related to cognitive ability, little is known about the neural correlates of normal variation of telorism within the general population. We derived measures of orbital telorism from cranial magnetic resonance imaging (MRI) by calculating the distance between the eyeball center of gravity in two population-based datasets (N=5,653, N=29,824, Mean age 64.66, 63.75 years). This measure was found to be related grey matter tissue density within numerous regions of the brain, including, but surprisingly not limited to, the frontal regions, in both positive and negative directions. Additionally, telorism was related to several cognitive functions, such as Perdue Pegboard test (Beta, P-value, (CI95%) −0.02, 1.63^×10-7^(−0.03;-0.01)) and fluid intelligence (0.02, 4.75^×10-06^ (0.01:0.02)), with some relationships driven by individuals with a smaller orbital telorism. This is reflective of the higher prevalence of hypo-telorism in developmental disorders, specifically those that accompany lower cognitive lower functioning. This study suggests, despite previous links only made in clinical extremes, that orbital telorism holds some relation to structural brain development and cognitive function in the general population. This relationship is likely driven by shared developmental periods.

## INTRODUCTION

Orbital telorism, the interocular distance, is a distinctive craniofacial trait which also serves as a clinically informative measure in its extremes. Orbital telorism is mainly determined in the embryonic period, driven by processes determining not only orbital telorism but other cranial features as well. As a result, extremes in orbital telorism often co-occur with other minor physical anomalies (MPAs). Pathways involved in embryonic development are commonly affected by genetic syndromes, some of which have hypo- or hypertelorism as one of the characteristic features ^1,2^.

In addition to MPAs, learning difficulties and cognitive limitations often accompany hypo/hyper telorism. For example, hypotelorism is more prevalent in autism, but, notably this is more often observed in lower functioning autistic individuals ^3,4^ Similar rates of hypotelorism as that in autism are also observed in those with developmental disabilities, suggesting it is not specific to autism but more common in persons with syndromes encompassing lower cognitive functioning and/or more severe developmental delay ^3,5^. Similarly, a higher prevalence of MPAs is reported within neurodevelopmental disorders ^6,7^. This could suggest that differences in brain development is also evident, with similar mechanisms at play. The relationship between telorism, brain structure and cognition along the whole spectrum of function within the general population is, however, not known.

To further elucidate the relationship between orbital telorism, structural brain features, and cognitive performance, we performed investigations into orbital telorism in relation to brain- and skull-structure, and cognition. We investigated the relationship between orbital telorism, structural brain and skull measurements, and cognition.

## METHODS

### Study Populations

#### UK Biobank

The UK BioBank (UKB) is a population-based prospective cohort with data collected at 23 research centres within the UK ^8, 9^. At the time of analysis 32,115 persons from the total sample had available T1 MRI scans of which 30,191 were usable for segmentation. Of 30,191 participants with usable MRI scans 367 failed registration and QC. The mean age was 63.75 years (SD 7.5) and women made up 52.2% of the sample. Ethnicity was defined as Caucasian (0), and other (1) including African, Asian, mixed and other classifications. The sample was predominantly Caucasian (97.1%).

#### Rotterdam Study

The Rotterdam Study (RS) is a population-based prospective cohort which started in 1990 in Rotterdam, the Netherlands ^10^. In 2006, magnetic resonance imaging (MRI) was included in the study protocol ^11^. At the time of analysis, 5642 had available T2 MRI scans. Of 5642 participants with usable MRI scans, 107 failed registration and QC. The mean age was 64.96 years (SD 10.9) and women made up 55% of the sample. Ethnicity was defined as Caucasian (0), and other (1) including African, Asian, mixed classifications. The sample was predominantly Caucasian (97.3%).

### Neuroimage acquisition

#### UK Biobank

Within the UK Biobank T2 scans were unavailable thus T1 weighted images were used. These T1-weighted images were performed on 3T Siemens Skyra and a 32-channel head coil across three imaging centers^8^. The images were collected using a sagittal 3D magnetization prepared-rapid gradient echo sequence with slice thickness of 1mm ^8, 12^.

#### Rotterdam Study

Within the Rotterdam Study T1-weighted and T2-weighted images were available. These were performed using a 1.5T MRI scanner (GE Signa Excite, General Electric Healthcare, Milwaukee, USA) with an 8-channel head coil ^11^. The T1 and T2 images were collected using a 3D GRE and 2D sequences respectively with slice thickness’ of 1mm and 1.6mm, respectively. Image acquisition parameters have been previously described in detail ^11^. A subset of the participants (n=85) were rescanned within 2-6 weeks after the first scan for reproducibility analysis.

### Neuroimage processing

#### Orbital telorism and skull measurements

Image processing for orbital telorism was performed on T2-weighted sequences, and if not available T1-weighted sequences were used. In short, advanced normalization tools (ANTs) registration was performed between the native and standard space (MNI^13,14^). Then, the eyeball mask manually derived from the MNI template was registered to the native space using the metrics from the initial registration. The centre of gravity of the eyeballs was used to calculate the Euclidian distance between the voxel coordinates in native space (Figure 1). The distances were manually checked if the distances, eye mask coordinates or mean voxel intensity, were more than 2.5 standard deviations away from the mean (<1% removed due to quality check). Due to a higher prevalence of hypo-telorism in neurodevelopmental disorders, a non-linear relationship is possible within the analyses, thus within this non-clinical population we also created sex-specific Hypo- and Hyper-telorism categories using -/+ 2 standard deviation cut offs within each sample.

**Figure 1.**
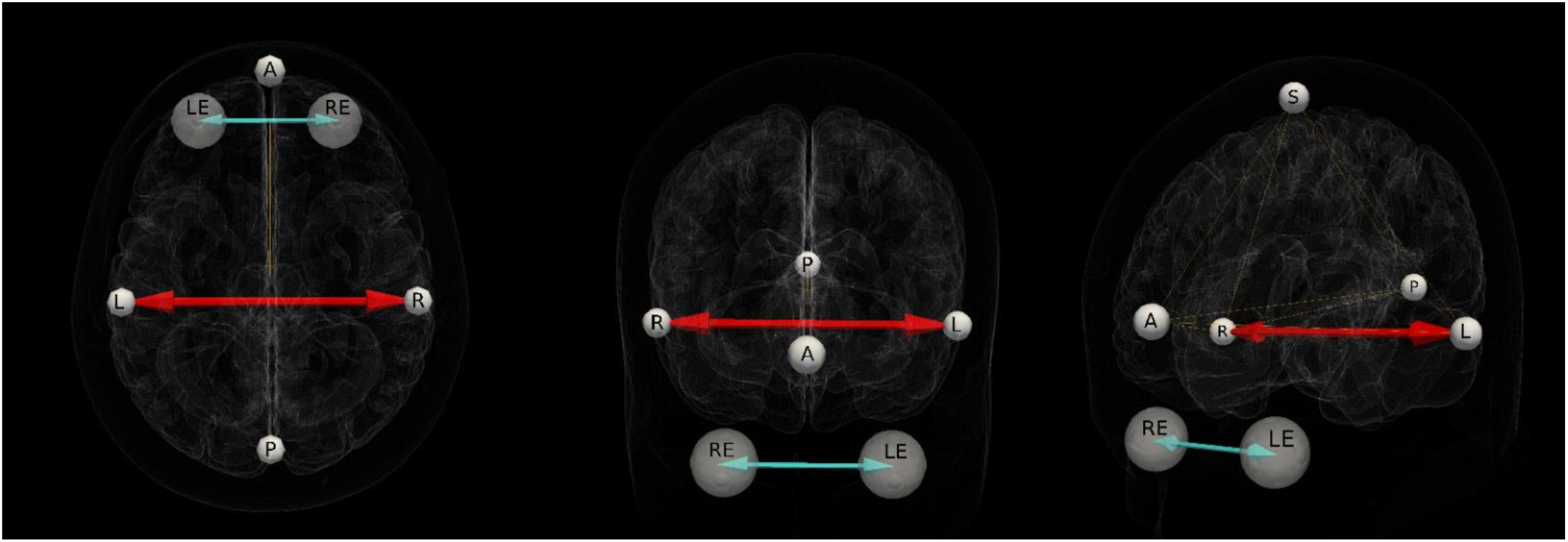
Orbital Telorism and Skull points: Methods. Measurements of orbital telorism and fiducial points. RE: right eye, LE: left eye. Skull points-R: right, L: left, A: anterior, PL: posterior, S: superior. Turquoise depicting orbital telorism (measurement from center of gravity of LE and RE). Red depicting head width (measurement from point L to R).

Similarly, a mask containing fiducial points on the inner skull of the MNI template was also registered to native space to extract skull measurements. In order to properly position the inner skull fiducial points, we identified points at the inner skull surface at the region of the temporal bone on a reference computed tomography atlas co-registered to the MRI atlas file ^15^. Position of the fiducial points in the MNI atlas was [x,y,z]; Left [68.3103 −30 −19.70263], Right [−69.11019 −30.69375 −19.70263], Superior [0 −30 87], Anterior [0 73.91968 12.01424], and Posterior [0 −99.16595 −15.6385]. Template based measurements and fiducial point locations can be seen in Figure 1.

In a reproducibility set of 85 individuals from the Rotterdam study we assessed the intraclass correlation coefficient (ICC) for orbital telorism and fiducial points using repeat MRI. The orbital telorism has shown a good reproducibility in the RS reproducibility subset (N = 85), with intraclass correlation coefficients (ICC) of 0.991 (95% confidence interval (CI) 0.98:0.99). Skull measurements also showed a good reproductivity in the RS reproducibility subset (N=85) with intraclass correlation coefficients (ICC) of: Width, 0.982 (0.97;0.99), Right-Superior, 0.86 (0.79:0.91), Right-Anterior, 0.88 (0.82:0.92), Right-Posterior, 0.91 (0.86:0.94), Left-Superior, 0.88 (0.81:0.92, Left-Anterior, 0.90 (0.85:0.94), Left-Posterior, 0.90 (0.85:0.94), Superior-Anterior, 0.94 (0.91:0.96), Superior-Posterior, 0.85 (0.78:0.91) Anterior-Posterior,0.99 (0.98:0.99).

In addition, for a subset of RS individuals’ (N=316) measurements of interpupillary distance via eye examination was available. We compared these measurements with our MRI derived orbital telorism measurements since they should highly correlate. This showed a good correlation with the measures of orbital telorism (ICC 0.84 (95% CI 0.80:0.87)).

### Intracranial volume

#### UK Biobank

Grey matter Tissue segmentation for VBM and intracranial volume (ICV) was calculation was done through the FreeSurfer software *recon* segmentation method within the UKB ^16^.

#### Rotterdam Study

Within RS MR images were processed into grey matter, cerebrospinal fluid, normal appearing white matter and white matter lesions using an automated processing algorithm, as described before ^11^, these were summed for calculation of ICV.

### Voxel based morphometry

To investigate the relation between orbital telorism and grey matter tissue at a focal level to address subtle differences voxel-based morphometry (VBM) was used. VBM was performed according to a previous protocol ^17^. Briefly, after tissue segmentation using the above-mentioned processes FSL software was used for VBM data processing ^16^. All GM density maps were registered (non-linear registration) to the ICBM MNI152GM standard template (Montreal Neurological Institute ^14,18^) with a 1×1×1mm^3^ resolution. Spatial modulation as used to avoid differences in absolute GM volume due to registration via multiplying voxel density values by the Jacobian determinates estimated during spatial normalization. Lastly, images were smoothed using a 3mm isotropic Gaussian kernel.

### Measures of cognition

#### UK Biobank

Cognition was measured using screen based cognitive tests within UKB ^19^. Pairs matching task consists of individuals asked to memorise the position of as many matching pairs of cards as possible. The participant is then asked to recall these pairs when the cards are faced down. The number of incorrect matches were used. The data is skewed therefore is log transformed before use in the analysis. The fluid intelligence test measures the capacity to solve problems requiring logic and reasoning ability, this was tested via an individual solving as many questions as possible from this 13-verbal logic/reasoning type multiple choice questions within 2 minutes. Number of correct answers were used within the analysis. Reaction time was assessed based on 12 rounds of the card game ‘Snap’. The participant is shown 2 cards at the same time and is to press a button as quickly as possible if these cards are the same. The score used was mean response time across 4 trials, of 12, that contained matching pairs. Symbol digit switching task, was used to assess cognitive switching via presentation of a grid linking symbols to single-digit numbers and a second grid containing only the symbols. Using the first grid as a key the individuals has to indicate the numbers attached to the symbols in the second grid. For analysis the number of correct symbol digit matches were used. For assessment of visual attention, the trail making test was used to provide information on visual search, mental flexibility, processing speed and executive function. In this task individuals are presented with sets of digits and letters in circles on a screen. The task is to connect the ascending sequence of numbers (Trail A), or alternating letters and numbers (Trail B). Time taken was used (log-transformed) for the analysis. Education was measured via years of education.

#### Rotterdam Study

Cognitive tests within the Rotterdam Study were performed during a centre visit ^20^. The Letter-digit substitution task (LDST) required individuals to make as many letter-digit combinations within 60 seconds as possible. The Perdue pegboard test is used to assess bilateral fine manual dexterity. This is done using a pegboard with two rows of 25 holes, individuals have to place as many pins as possible in a prescribed order within 30 seconds. Trials using both hands were used within this analysis with a summary score calculated over 3 attempts. Stroop tests, abbreviated form, were used consisting of reading, colour naming and interference tests consisting of naming printed words, naming printed colours and naming the colour in which a colour-name is printed, respectively. Lower scores on Stroop tests indicate a better score. To asses memory the world learning tests (15-WLT), based on Rey’s auditory recall of words, was used. These included individuals being presented with 15 words and asked to recall as many as possible immediately after (immediate test), and also 15 minutes later (delayed test). Additionally, recognition memory was assessed via these words being presented with a further 30 words with participants having to identify the original words. Correct scores were used for the analysis. Lastly word fluency was assessed using the word fluency test (WFT), asking participants to name as many animals as possible within 60 seconds. Education was recorded as years in education.

### Statistical analysis

Linear regressions were fitted to assess the relation between orbital telorism and skull measurements. For skull analysis, models were adjusted for age, sex, and ethnicity, with Model II additionally adjusted for ICV. Within VBM analysis, linear regression models were fitted with voxel values of GM modulation density as the dependent variable and orbital telorism value/group membership (hypo-hyper telorism) as independent variables, adjusting for age, sex, ethnicity, and head width. Previous permutation testing was carried out to compute a corrected P value of 2.99×10^−07^ for VBM analysis ^21^. The results from RSS and UKB were meta-analysed using a fixed effects meta-analysis for each voxel. Additionally, linear regressions were fitted to assess the relation between orbital telorism and hyper- and hypo-telorism classifications, and cognition. For this cognition analysis an adjusted threshold of 0.00051 was used based on permutation analysis. Models were adjusted for: Model 1. age, sex, and ethnicity, Model II plus education (years), and Model III plus head width.

## RESULTS

### Descriptive characteristics

We included 35,459 middle-aged and elderly participants the UK Biobank (UKB) (29,824) and from the Rotterdam Study (RS) (5,535). Descriptive information about these studies, including population characteristics, cognitive tests are presented in Table S1-3. When characterizing hypo- and hyper-telorism via sex specific −/+ 2 SD this resulted 1.9% and 2.8% for UKB, and in 1.6% and 3.3% for RSS for hypo- and hyper-telorism groups respectively.

### Skull relations to orbital telorism

Subsequently, we aimed to determine the relationship between skull measurements and orbital telorism. Orbital telorism was significantly related to all measures of the skull that were extracted in a positive direction, however many relationships attenuated after adjustment for ICV within the Rotterdam study (Figure 2, Table S4). This attenuation was observed marginally within the UK Biobank sample however the relationships remained significant.

**Figure 2.**
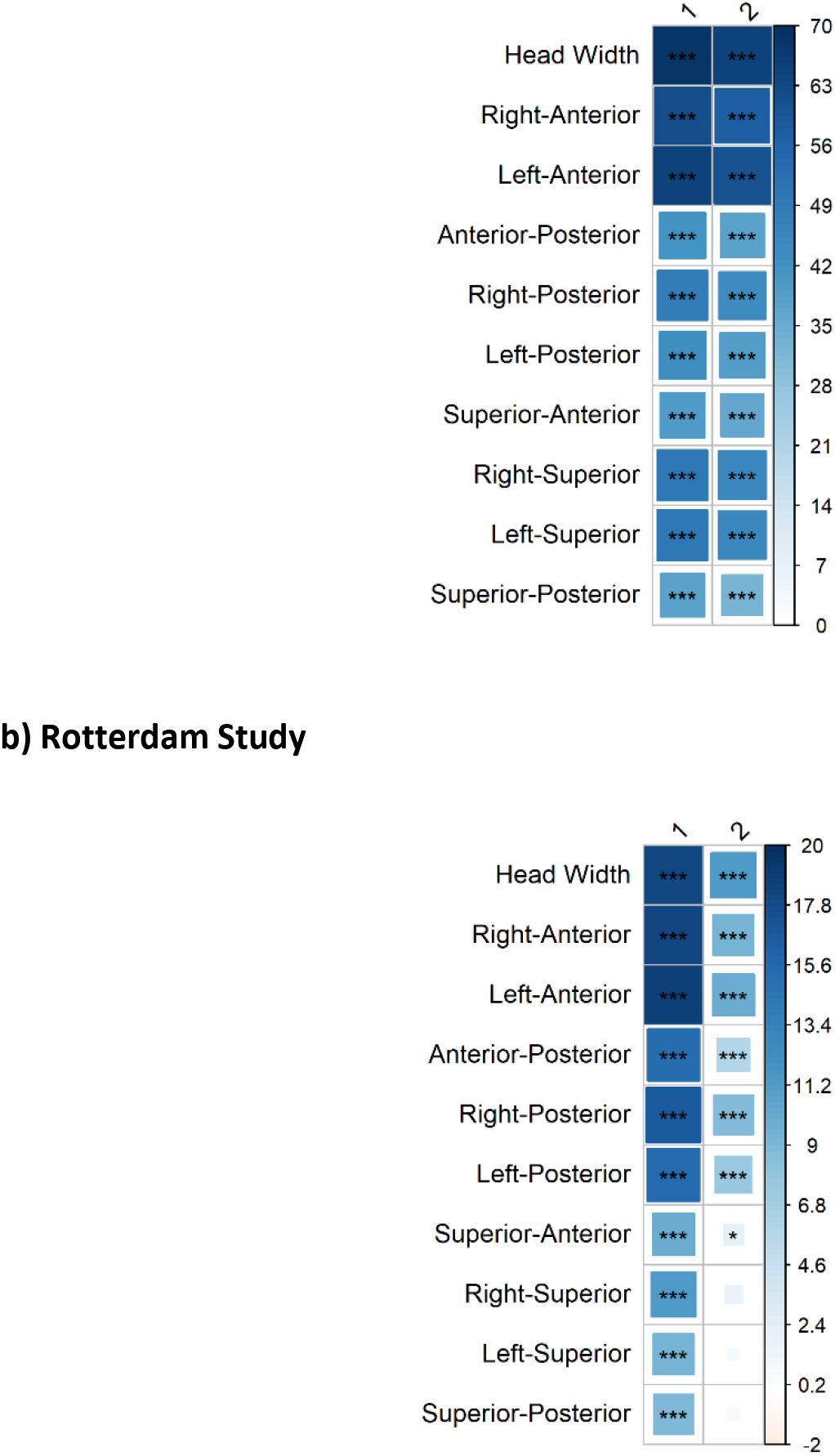
Association between Orbital Telorism distance and skull measurements. Colour intensity and number scale represents t value. Model 1; age, sex, ethnicity (white/non) Model 2: additionally, adjusted for ICV. P values: ***<0.0051, **, P<0.01, *<0.05. NOTE: Range of t values differ for panel a (−2:20) and b (0:70).

### VBM

When the two samples VBM results were meta-analyzed (Figure 3, Table S5) the strongest positive associations of orbital telorism with grey matter voxels (adjusting for age, sex and ethnicity) were the medial orbital gyrus (left; 4816, 4.28×10^−161^, right; 3726, 1.13×10^−220^), straight gyrus (left; 4544, 1.92×10^−188^, right, 5248, 1.34×10^−205^) and the subgenual anterior cingulate gyrus (left; 1144, 7.7×10^−114^, right; 1337, 2.54×10-^195^). The strongest primarily negative associations were found with the amygdala (left; 1204, 4.07×10^−115^, right, 1307, 5.11×10^−91^). Regions that displayed strong positive and negative relations were (positive and negative) right superior frontal gyrus (8070 and 2469, 4.79×10^−212^) and the middle frontal gyrus (left; 10815 and 1351, 3.14×10^−115^, right; 7303 and 1017, 1.45×10^−204^).

**Figure 3.**
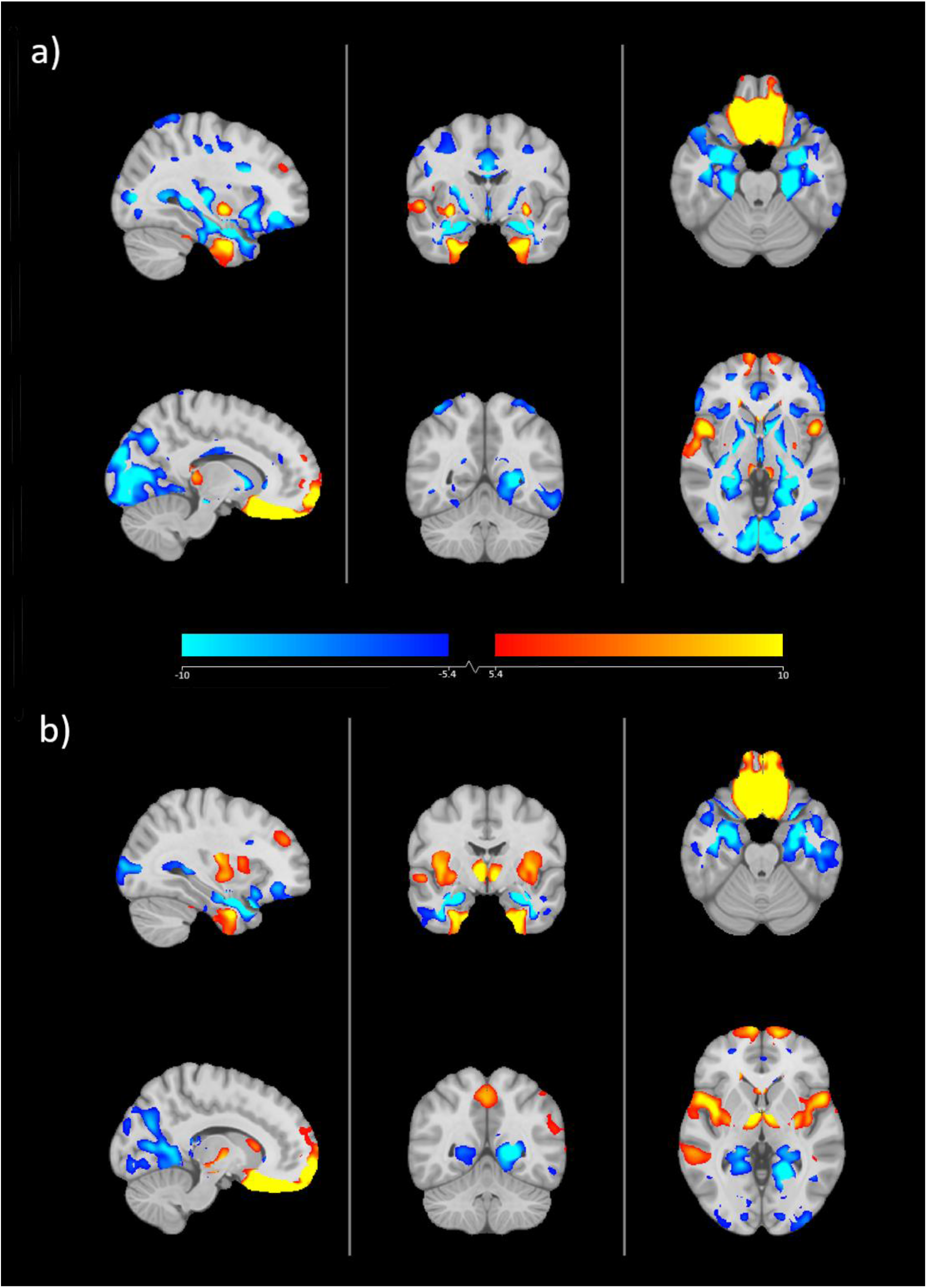
Telorism VBM Meta-analysis results over RSS and UKB. Heat map and colour bar represent t values to the significance threshold. Red indicates positive direction, blue indicates negative directions. Model I (a) adjusted for age, sex, ethnicity, Model II (b) additionally adjusted for head width.

When additionally adjusting for head-width, similar relations were seen, albeit with generally smaller voxel clusters and more significant relations (Figure 3, Table S5). However, adjusting for head-width did appear to attenuate many negative clusters. For example, the strongest positive associations of orbital telorism with grey matter voxels were found in medial orbital gyrus (left; 5561, 4.3×10^−180^, right; 4937, 3.4×10^−275^), straight gyrus (left; 4819, 2.2×10^−303^, right; 5640, 7.5×10^−271^) and the subgenual anterior cingulate gyrus (left; 1160, 2.4×10^−121^, right;1309, 3.7×10^−234^). The strongest negative associations of orbital telorism with grey matter voxels were found in the Amygdala (left 1091, 1.7×10^−52^, right, 1231, 3.0×10-53) and medial and inferior temporal gyri (left; 9633, 1.8×10^−56^, right; 7074, 9.9×10^−52^).Regions that displayed strong positive and negative relations were superior frontal gyrus (positive and negative) (left; 5689 and 319, 2.6×10^−116^, right; 8001 and 1107, 1.4×10^−256^), the middle frontal gyrus (left: 2849 and 1368, 2.8×10^−122^, right; 2002 and 1424, 2.4×10^−242^), anterior orbital gyrus (left; 2045 and 1357, 1.3×10^−87^, right; 1745 and 1115, 9.9×10^−108^) and the left insula (5712 and 1204, 2.9×10^−50^).

In the hypo- and hyper-telorism VBM, similar clusters are seen (Model II) with the strongest positive associations reported in the straight gyrus and the medial orbital gyrus, and negative associations within the medial and inferior temporal gyri (Figure FS1a-b). Negative associations that were seen between continuous orbital telorism measures and posterior temporal lobe and the superior parietal gyrus were also seen within the hyper telorism analysis, however this cluster was much smaller or non-existent in the hypotelorism analysis. A full break down of the meta-analysis and sample specific results can be found in Table S5-6.

### Cognition

Orbital telorism was found to be associated to measurements of manual dexterity, fluid intelligence, fluency and visual attention and task switching.

Within the UK Biobank higher scores of fluid intelligence was related to increasing orbital telorism measurements in Model III (0.03 (0.01:0.02)) (Figure 4a), however this was driven by the lower measurements of orbital telorism as seen when classifying the groups (Hypotelorism, M I; −0.16 (−0. 29:-0.02)) which also attenuated when adjusting for head width (Figure 5b). Furthermore, when classified into subgroups hypotelorism was also related to worse functioning on the trail making test, subtest B (Model II, 0.15 (0.06:0.24)(Figure 4b) this relationship attenuated to below the adjusted significance threshold after further adjustment for head width (M III) (0.11 (0.02:0.20) (P<0.05). Full results can be seen in Tables S7-10.

**Figure 4.**
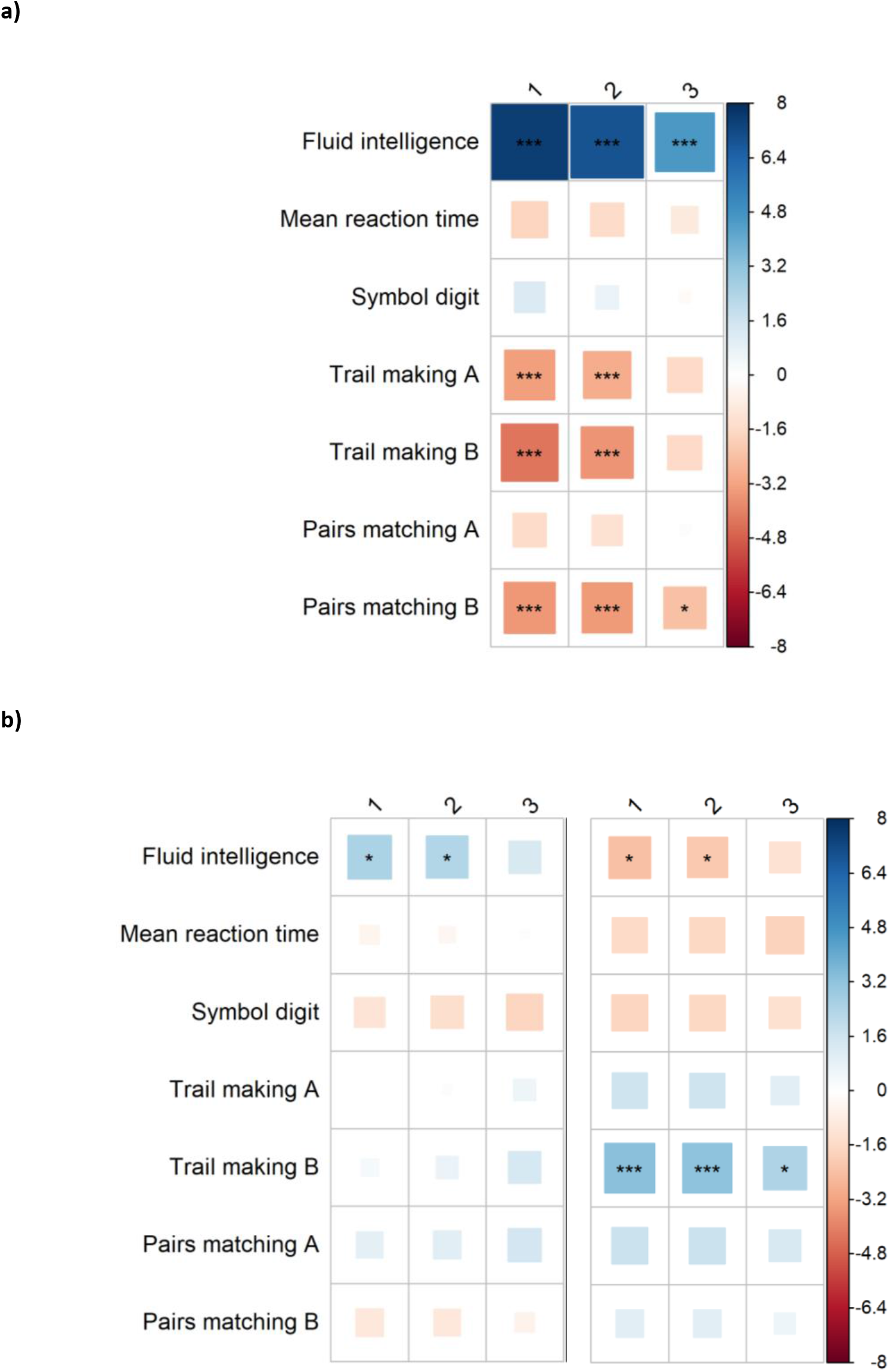
Association between Orbital Telorism distance and cognitive tests in UKB. Figure depicts full sample (a), grouped analysis (b-Left, Hyper-telorism, Right, Hypo-telorism). Model I Age + sex + ethnicity, Model II + Education, Model III + Head width. Reaction time and trail making tasks are in the inverse direction where a higher score indicates a worse function. P values: ***<0.0051, **, P<0.01, *<0.05. Trail making tasks are log transformed (ln).

**Figure 5.**
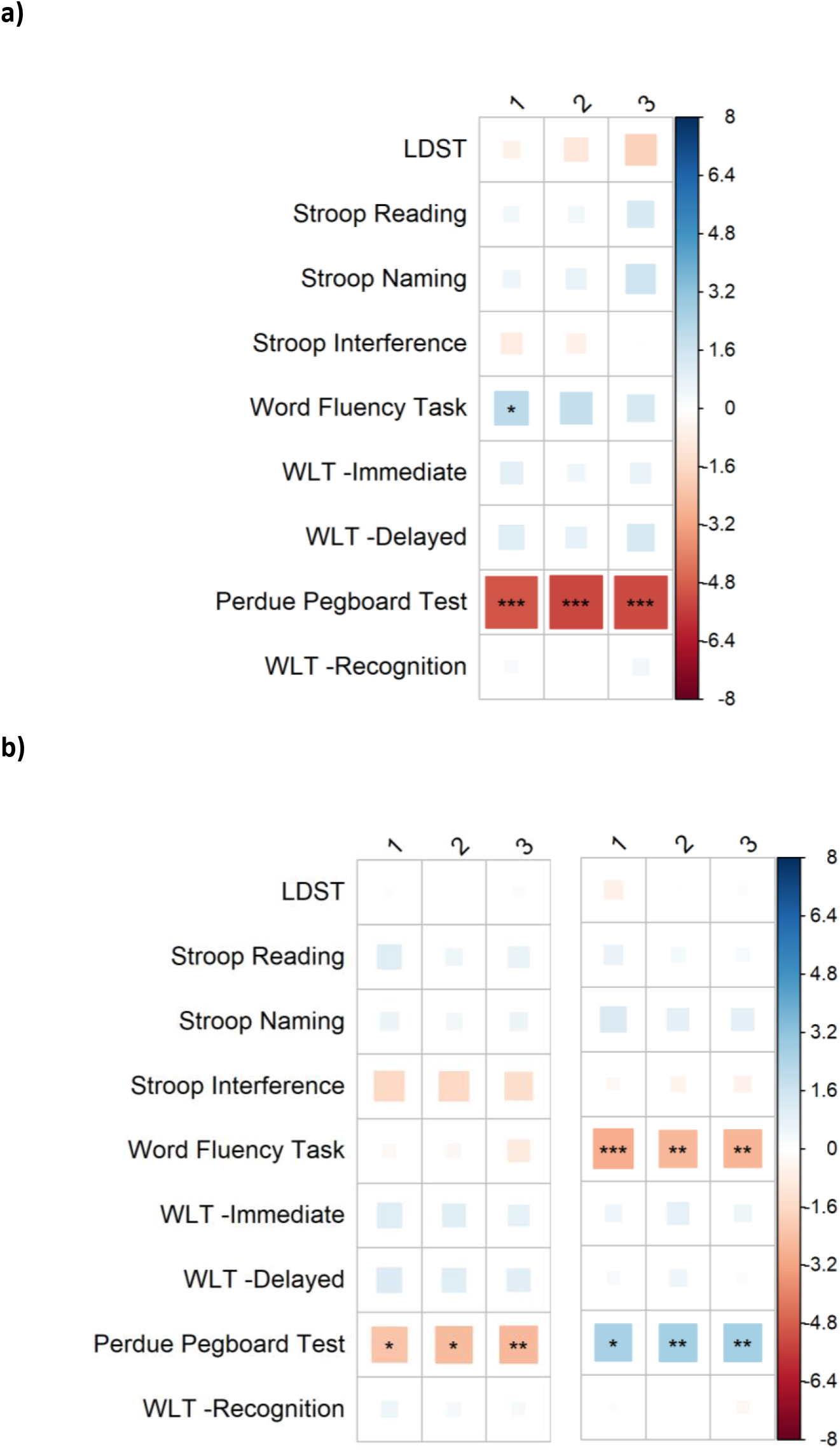
Association between Orbital Telorism distance and cognitive tests in RSS. Figure depicts full sample (a), grouped analysis (b-Left, Hyper-telorism, Right, Hypo-telorism). Model I Age + sex + ethnicity, Model II + Education, Model III + Head width. Stroop relations are in the inverse direction where a higher score indicates a worse function. P values: ***<0.0051, **, P<0.01, *<0.05. LDST; Letter digit substitution task, WLT; Word learning task.

Within the Rotterdam Study, relations between continuous measures of orbital telorism and the Perdue Pegboard test were found with higher telorism measurements related to lower test scores (Beta (CI95%)) (Model III, −0.02 (−0.03:-0.01) (Figure 5a). Additionally, when classifying the measurements into hyper- and hypo-telorism, classification of hypotelorism was associated to lower scores of word fluency (Model I, −0.32(−0.53:-0.11))(Figure 5b). However this attenuated after adjusting for education and head width(Model III, −0.29 (−0.50:-0.08))(P<0.05). A similar relationship was not observed in those within the hypertelorism group (Model I, −0.02 (−0.17:0.12)).

## DISCUSSION

Due to the higher prevalence of clinical extremes of orbital telorism within those with developmental disabilities, we studied if orbital telorism, brain structure and cognition displayed a relationship within the general population. We identified skull measurements, and cortical and subcortical structures related to orbital telorism beyond the regions located closely to the orbits. We also found that orbital telorism is related to performance in multiple cognitive tests, such as fluid intelligence, independent of head width, in a large population-based sample.

Skull structure is a highly conservative and complex structure. We anticipated that skull shape measurements derived from fiducial points located in the inner surface of the skull will have close relationship to the orbital telorism. This was indeed the case, with the strongest relationships unsurprisingly found with the anterior part of the skull. Across the skull measurements, relationships to orbital telorism were not uniform, with the other measurements, such as those connected to the superior part of the skull, found to be much weaker.

The mechanisms that control gradual brain and skull growth during early development are carefully timed and controlled through growth factors^22^. Skull growth dynamics are precise enough that the orbital telorism has been proposed as a marker of foetal age^23^. Growth can be altered by a multitude of environmental factors, for example through fibroblast growth factor 2 interacting with the family of glypicans belonging to cell surface heparan sulfate proteoglycans^24^, and modulated by genetic influence, like the modulators of Sonic Hedgehog pathway^25^ leading to final interocular distance in the course of development. Additionally, presence of certain alkaloids in the diet during early brain development has been found to alter the signalling and influence the growth of the brain and interocular distance^26^. These shared mechanisms influencing brain and skull growth result in the timing, dynamic and regional variability of skull growth being tightly linked to brain development.

Within the current study, we found orbital telorism and regional volume of brain structures to be related even after controlling for head width (as a proxy for skull development), which might indicate that exposure to FGF2 signalling and Shh pathway has differential influence over early development of subcortical regions. Unsurprisingly, the brain regions located closely to the orbits were highly positively related to the orbital telorism distance. That indicates the dependence of brain structure volume and the shape on the intracranial space is possibly defined by anterior regions of the cranial cavity. We also identified several negative relationships between the interorbital distance and areas of subcortical and posterior brain regions. Examples of these negative relationships are between orbital telorism and the amygdala and the hippocampus. This suggests that a shorter orbital telorism distance is related to a larger amygdala and hippocampus. Whilst research is inconsistent regarding amygdala and hippocampal volume in those with neurodevelopmental disorders ^27–31^, a larger amygdala has previously been noted to be associated with neurological and neuropsychiatric disorders, such as major depressive disorder^32,33^. Furthermore, the amygdala and hippocampus are suggested to be functionally abnormal and have an abnormal structure in autism and other neurodevelopmental disorders, disorders that additionally have a higher prevalence of orbital hypotelorism^34,35^. Possibly explanations for these negative findings are that these structures develop during periods also important for skull changes and the movement of the orbital bones where the angle between the orbits decreases. Limited and restricted growth and reduced perfusion could impact specific regions within specific developmental periods. Additionally, negative relationships may also be present due to compensation from larger previously developed areas. Whilst fetal MRI is still in relatively early stages, understanding the co-occurrence of normal and abnormal development of brain structures and skull formation would be deeply interesting to further understand pathways involved.

The thalamus also displayed negative relations, but this was also accompanied by positive clusters within the reasonably large subcortical structure. This would be interesting to further investigate a more functionally parcellated segmentation of the thalamus to understand the differences in these clusters and how this relates to developmental pathways. The corpus callosum also displayed both negative and positive clusters within the VBM analysis. This vital structure is involved within the visual and motor system, interhemispheric connectivity and is also involved in regulation of emotional and cognitive functions^36–39^. Previous studies identified abnormal morphology of the corpus callosum in neuropsychiatric disorders such as OCD ^40–43^. We hypothesize that the genes responsible for early brain development might play a significant role in neuropsychiatric disorders. More research investigating this relationship within clinical samples and into possible biological pathways that could be involved in development of these structures is needed.

Since we identified specific brain regions associated with orbital telorism, we additionally were interested to see whether this trait was also related to cognitive function. Despite minor physical anomalies, such as clinical classifications of hypotelorism, being reported to have higher prevalence in neurodevelopmental disorders, the relationship between normal variation in orbital telorism and cognition was not yet studied. The current study finds links between normal variation of orbital telorism and cognition within the normal population. One drawback, however, of the current study is the lack of correspondence between the cognitive tasks within the two included samples, hindering the ability to compare and meta-analyse. Despite this, they do give insight into a wider range of cognitive functions than either alone.

The study found multiple relations between orbital telorism and cognitive ability, with some observations appearing to be driven by the hypotelorism end of the spectrum. Firstly, an association between motor function and orbital telorism was seen. The association between the Purdue pegboard test performance and orbital telorism within the Rotterdam Study sample was initially surprising. This test is mainly used to investigate manual dexterity; however, it is also possible other factors may be driving this relationship. The task involves spatial perception to be able to efficiently move the pieces into the correct locations. This relation could be driven by the worsening of perception as the orbital telorism distance increases for this task. We also think that the relationship might be linked to development of motor coordination and interhemispheric communication and visual processing speed^36,37,44^. This is suggested due to the relationship of the corpus callosum with visual and motor tasks previously indicated, a structure also associated to orbital telorism within the current VBM analysis.

When investigating the extremes of the sample, the hypotelorism group was also found to have worse scores on word fluency, a relationship not seen in the continuous analysis. This test is also related to worse function in individuals with neurodevelopmental disorders, further underlying the link between neurodevelopmental dysfunction and hypotelorism, even in the extremes of normal within the general population ^6,7^.

Further investigations into the relationship between orbital telorism and cognition within the UK Biobank provide further support for the relationship between orbital telorism and cognition. The main results from this investigation indicate that orbital telorism is positively related to fluid intelligence. Previous research has indicated that the prefrontal and parietal regions are important for fluid intelligence, these regions were also implicated within the orbital telorism VBM and could be directly linked to the development of these brain regions and skull morphology. Furthermore, the relationship to a larger orbital telorism and fluid intelligence is further supported by the prevalence of hypotelorism being higher in low-functioning individuals with autism and those with developmental delay^3,5^. These results suggest that this relationship between orbital telorism, possibly in this aspect a proxy of skull and brain development, and cognition can also be found within the general population, albeit on a smaller, and less clinically relevant, scale.

Telorism is reported to differ across different ethnic groups ^45,46^. Genetic underpinnings of skull and facial morphology have been studied providing the grounding for these differences. Despite this, the current study does not include a wildly diverse sample demographic, with both samples from European regions, with a predominantly white background, thus limiting the generalizability of the current results. Despite this the method can be easily used within further samples in the future, and also possibly adapted to be applied to other modalities such as CT scans for wider use.

The current study is the first comprehensive investigation into the neuronal correlates of orbital telorism. In addition to this, the study’s strengths lie in its large sample and use of the best available robust method of extracting orbital telorism from structural imaging given the lack of face photograph or calibrated face scan. Additionally, the methods used are based on open-source software and thus can be replicated on various other datasets that include face on the MRI. For example, a lot of international samples include MRI thus this can therefore be widely applied on existing data without the need to create new facial photographs. Despite this, orbital telorism derived from MRI is not the gold standard for this area. However, the method was reported to show good correlation with manual measurements. Furthermore, MRI has the additional quality of not being affected by amblyopia, thus focusing on the measurement of the centre of the orbit and not the pupil.

In conclusion, this study provides various insights into the neural correlate relations to orbital telorism. The study highlights positive, but also negative relations for orbital telorism with subcortical brain structures and cognition in the general population that could be implicated within neurodevelopmental disorders. Further studies into clinical samples, such as neurodevelopmental samples, and more diverse samples, is needed to further investigate the relations between orbital telorism, cortical and subcortical brain regions and cognition and the use of orbital telorism as a useful biomarker.

We declare no competing interests.

## Supplement

### Tables

S1. Demographics

S2. Ethnicity

S3. Cognition means: (a) RSS, (b) UKB

S4. Association between Orbital Telorism distance and skull measurements

S5. VBM results Meta-analysis: (a-b) Orbital telorism, (c-d) Hyper-telorism, (e-f) Hypo-telorism

S6. VBM results individual: (a-f) UKB, (g-j) RSS

S7. Cognition results: RSS

S8. Cognition results: RSS, hyper/hypo-telorism

S9. Cognition results: UKB

S10. Cognition results: UKB, hyper/hypo-telorism

### Figures

FS1. VBM Meta-analysis hyper/hypo-telorism

