## Supplementary figures and images for "Neural correlates of orbital telorism"

### Supplemental Figure 1a

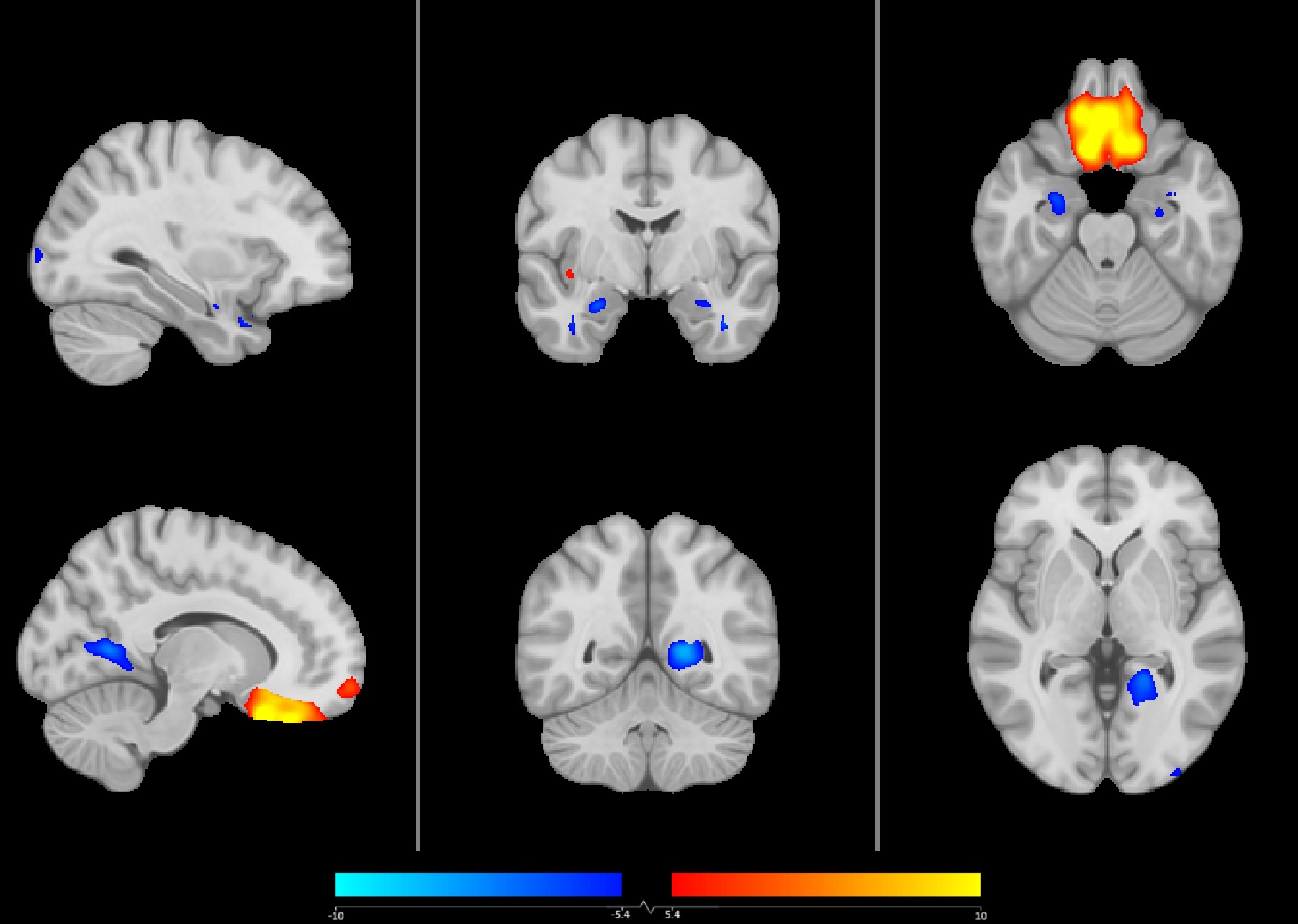

### Supplemental Figure 1b

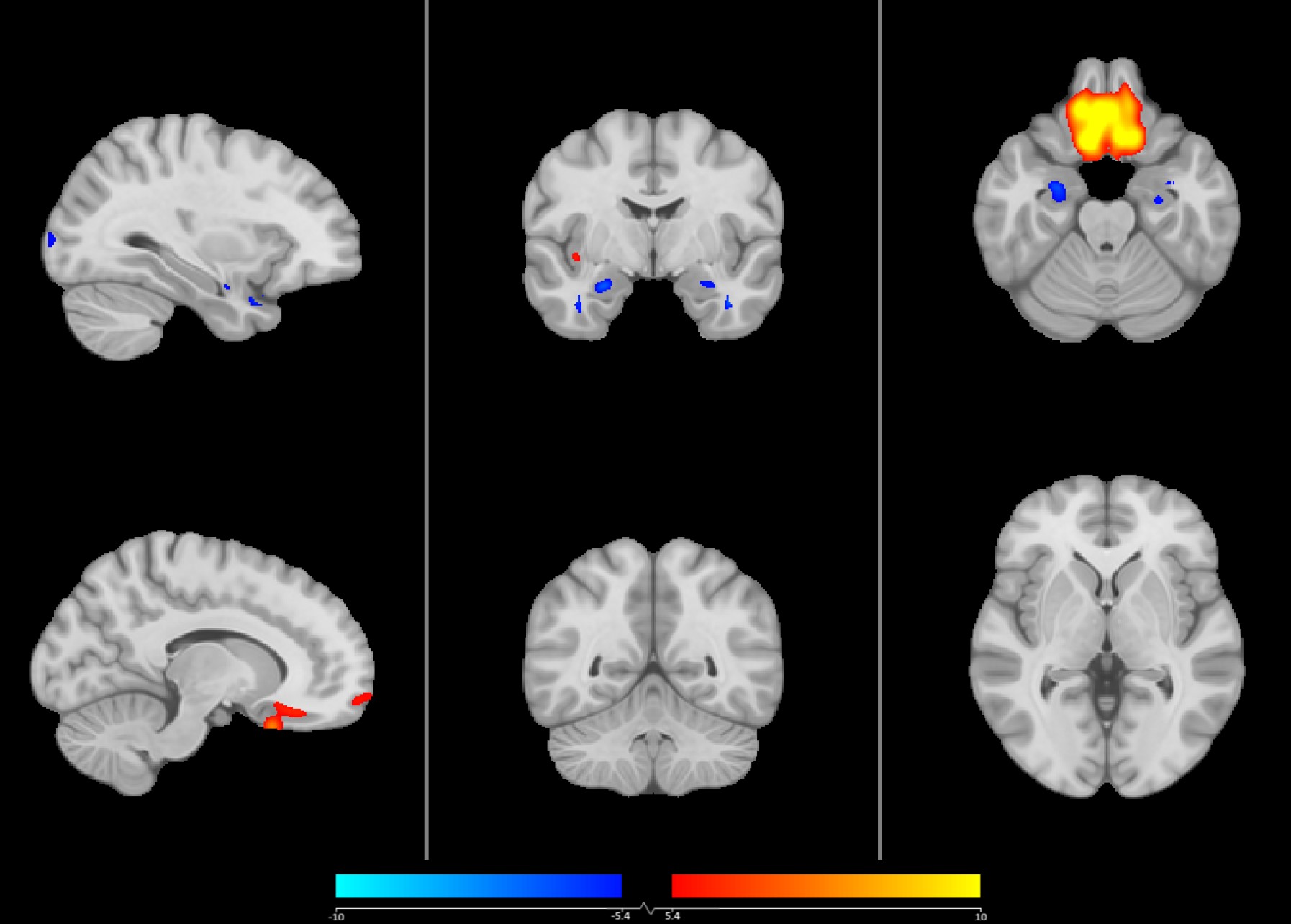
